# Inclusion of minor alleles improves catalogue-based prediction of fluoroquinolone resistance in *Mycobacterium tuberculosis*

**DOI:** 10.1101/2022.11.09.515757

**Authors:** Alice E. Brankin, Philip W. Fowler

**Affiliations:** Nuffield Department of Medicine, University of Oxford, UK; National Institute of Health Research Oxford Biomedical Research Centre, John Radcliffe Hospital, Oxford, UK

## Abstract

Fluoroquinolone resistance poses a threat to the successful treatment of tuberculosis. Whole genome sequencing (WGS), and the subsequent detection of catalogued resistance-associated mutations, offers an attractive solution to fluoroquinolone susceptibility testing. However, the bioinformatic pipelines used often mask the recognition of minor alleles which are implicated in fluoroquinolone resistance. Using the Comprehensive Resistance Prediction for Tuberculosis: an International Consortium’s (CRyPTIC) dataset of globally diverse WGS *Mycobacterium tuberculosis* isolates, with matched minimum inhibitory concentrations for two fluoroquinolone drugs, we show that detecting minor alleles increased the sensitivity of WGS for moxifloxacin resistance prediction from 85.4% to 94.0%, without significantly reducing specificity. We also found no correlation between the proportion of an *M. tuberculosis* population containing a resistance-conferring allele and the magnitude of resistance. Together our results highlight the importance of detecting minor resistance conferring alleles when using WGS, or indeed any sequencing-based approach, to diagnose fluoroquinolone resistance.

## Introduction

The fluoroquinolone antibiotics levofloxacin and moxifloxacin are recommended by the WHO for the treatment of *both* drug susceptible and multi-drug resistant (MDR) tuberculosis (TB)^1^. It is therefore imperative that fluroquinolone drug susceptibility testing (DST) is carried out quickly and accurately to treat patients and prevent the spread of resistant strains.

Whole genome sequencing (WGS) can rapidly identify resistance and susceptibility to several antitubercular drugs^2,3^. The WHO recommends that WGS results are interpreted using a catalogue of mutations associated with resistance compiled using over 38,000 *M. tuberculosis* isolates^4^. The sensitivity of this catalogue for identifying resistance was >90% for the first line drugs rifampicin and isoniazid in the dataset used to build it ^5^. However, the sensitivity to identifying levofloxacin and moxifloxacin resistance was lower, at 84.4% and 87.7% respectively^5^. Since fluoroquinolone resistance is well characterised and attributed to a small number of mutations in the *gyrA* and *gyrB* genes^6^ it is surprising that a larger proportion of fluoroquinolone resistance was not explained by the catalogue.

Mixed populations are common in *M. tuberculosis* infections and have been particularly implicated in fluoroquinolone resistance^8,9^; the prevalence of mixed populations containing minor resistance-conferring alleles is estimated at c. 10% of fluoroquinolone resistant isolates^10^. WGS bioinformatic pipelines often use filters and thresholds to reduce the effect of sequencing errors^7^, which unfortunately also preclude the detection of minor alleles.

Indeed the pipeline used in processing the samples for the WHO catalogue^4,5^ only identified a genetic variant if over 90% of reads at a genomic position supported its existence, effectively ignoring minor alleles.

Large matched WGS and phenotypic datasets, such as that compiled by the Comprehensive Resistance Prediction for Tuberculosis: an International Consortium (CRyPTIC)^11^, provide an opportunity to study the significance of minor alleles. In this paper we will investigate the extent to which catalogue-based fluoroquinolone resistance prediction is enhanced by including minor alleles containing known resistance-conferring mutations.

## Materials and Methods

We assume that a population is homogenous if 90% or more of the reads support a different nucleotide to the reference genome (i.e. fraction of read support, FRS ≥0.9), whilst a population is mixed if there are more than two reads but fewer than 90% supporting a genetic variant. Our rationale is that as the error rate of Illumina sequencing is <1%, if two or more reads support an alternative allele, it is highly unlikely that this is due to sequencing error.

*M. tuberculosis* complex isolates with an associated moxifloxacin or levofloxacin minimum inhibitory concentration (MIC) were obtained from the CRyPTIC FTP site (ftp://ftp.ebi.ac.uk/pub/databases/cryptic/release_june2022/). Isolates were discarded if the phenotypic measurement was annotated as ‘low quality’ (the three methods used to determine the MIC were not in agreement) or if they originated from a laboratory with a known quality control issue^11^. The genetic variants these isolates had in both the *gyrA* or *gyrB* genes, as detected by Illumina sequencing and the CRyPTIC bioinformatic pipeline^11^, were also extracted from the dataset. In total, 9,128 isolates with WGS and minimum inhibitory concentration (MIC) data for levofloxacin and 8,138 isolates with WGS and MIC data for moxifloxacin were analysed.

The variant caller (Clockwork v0.8.3, https://github.com/iqbal-lab-org/clockwork) used by the CRyPTIC project was set up conservatively; a genetic variant had to have an FRS ≥90% for it to be identified, with all other potential variants being screened out^11^. We therefore directly parsed the variant call format files of all samples looking for isolates that had evidence of minor alleles in *gyrA* or *gyrB*. The FRS for each putative genetic variant was extracted and any resultant amino acid changes identified. Putative variants present at a position which failed the Minos^12^ minimum sequencing depth filter were not included, and hence assumed wild-type for analyses. The minimum genotype confidence percentile was not used to exclude any variants because this filter is itself partially dependent on the FRS^12^. Finally, we assumed that genetic variants in the WHO resistance catalogue annotated as either ‘resistance associated’ and ‘resistance associated – interim’ mutations both conferred resistance to the relevant drug.

## Results

CRyPTIC isolates were classified as resistant or susceptible depending on whether their minimum inhibitory concentration (Fig. 1a,b) lay above or below a published ECOFF ^13^. We then predicted genetically which of the isolates were resistant to levofloxacin and moxifloxacin assuming the populations were homogenous (all genetic variants supported by an FRS ≥0.9). This approach identified the levofloxacin and moxifloxacin resistant isolates with 83.1% and 85.4% sensitivity respectively, and over 90% specificity in both cases (Fig. 1c,d). In total, 16.9% of levofloxacin resistance and 14.6% of moxifloxacin resistance in this dataset is therefore not explainable by catalogue mutations seen at FRS ≥0.9.

**Figure 1.**
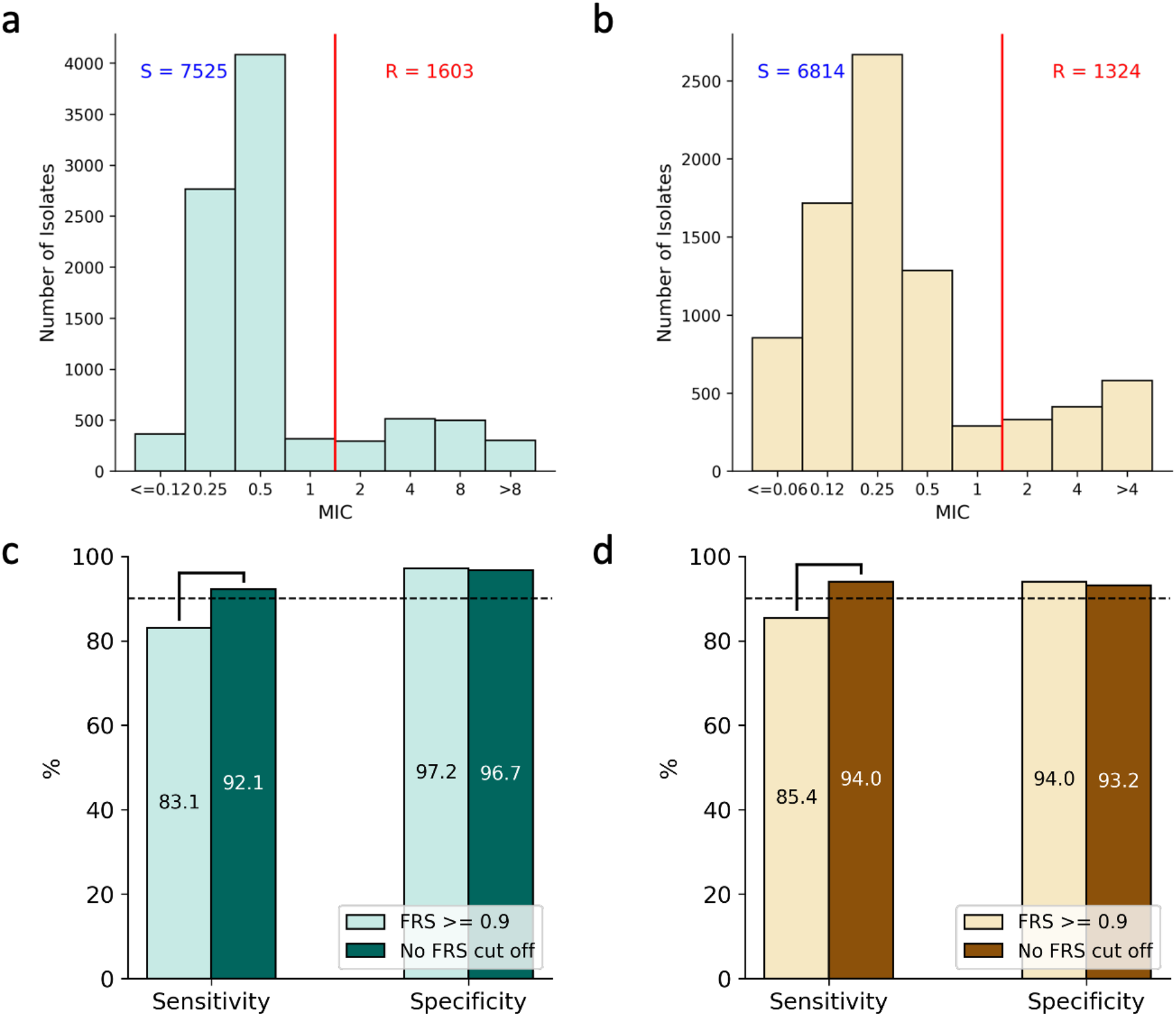
Distribution of *M. tuberculosis* isolates Minimum Inhibitory Concentrations to. **(a) levofloxacin and (b) moxifloxacin**. The red line indicates the previously proposed epidemiological cut off value (ECOFF)^13^ that was used to distinguish resistant (R) and susceptible (S) isolates. **(c) Sensitivity and (d) specificity of fluoroquinolone resistance prediction using WHO 2021 catalogue mutations with and without mixed alleles**. Brackets indicate a significant difference (z-test, p < 0.05).

Allowing minor alleles containing mutations in the WHO catalogue to contribute to the predictions by reducing the FRS threshold increased the sensitivity of the catalogue significantly, by 9.7% and 9.5% when identifying levofloxacin and moxifloxacin resistance, respectively (Fig. 1c,d). The specificity of the predictions, decreased slightly by 0.5% and 0.8% for levofloxacin and moxifloxacin, respectively, but these differences were not significant at *p* = 0.05. For this dataset, 7.9% of levofloxacin resistance and 6.0% of moxifloxacin resistance remains unexplained by the presence of catalogue mutations.

One might expect that if a resistance conferring allele is seen at higher FRS (i.e. is more prevalent within the mixed population) it will have a greater level of resistance. Hence we next examined whether the proportion of the population containing the minor resistance conferring allele (FRS) correlates with the magnitude of the fluoroquinolone MIC. Although all catalogue resistance-associated mutations were seen as part of a mixed population in at least one sample, we only considered the two most frequent mutations seen in fluoroquinolone resistant isolates, *gyrA* D94G and A90V. Pearson’s rank correlation coefficients confirm there is no correlation between FRS for the resistance-conferring allele and MIC to either levofloxacin or moxifloxacin in isolates with either *gyrA* D94G or A90V (Fig. 2a-d).

**Figure 2.**
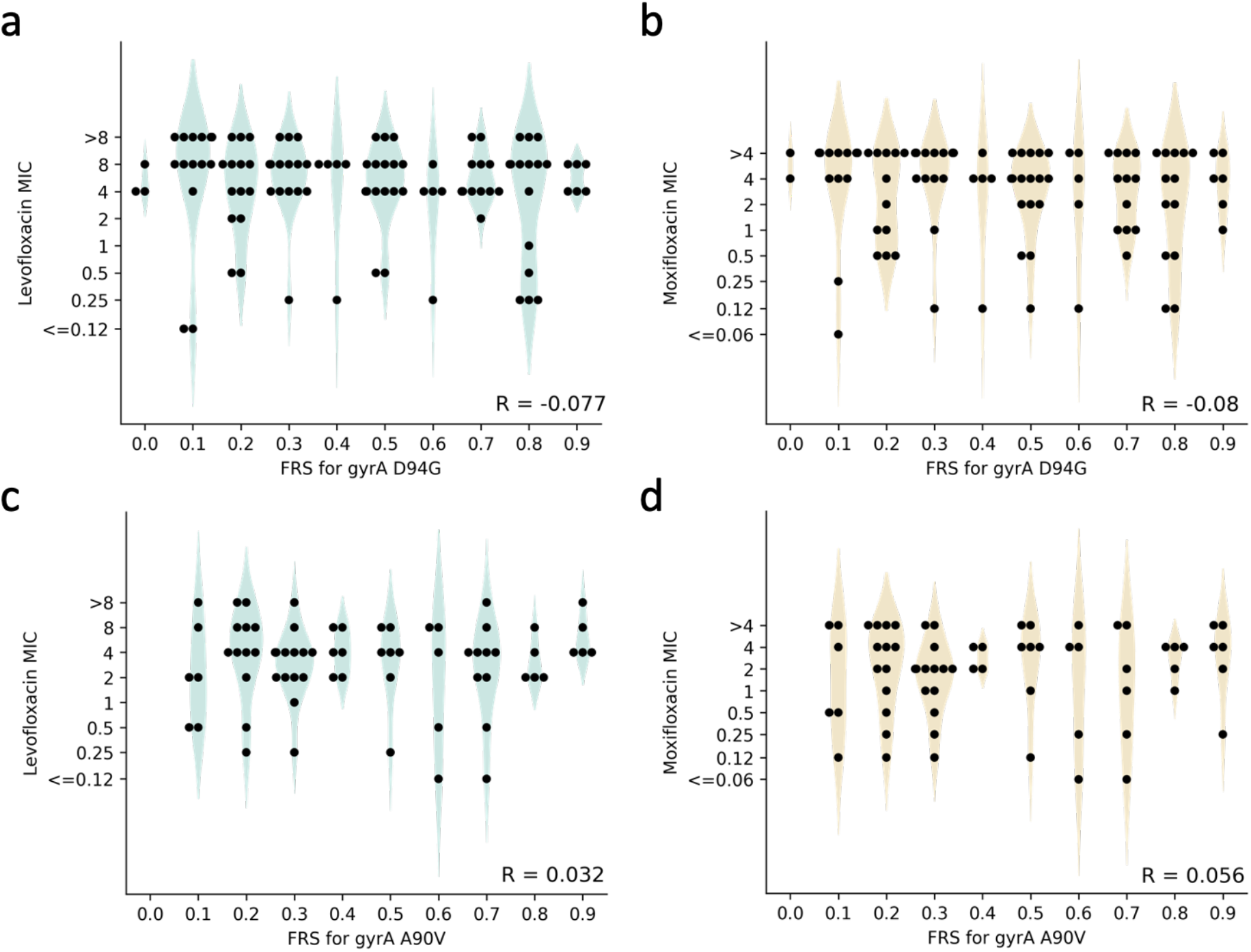
Distribution of FRS for *gyrA* D94G and the (a) levofloxacin and (b) moxifloxacin MIC of the *M. tuberculosis* isolate and the distribution of FRS for *gyrA* A90V and the (c) levofloxacin and (d) moxifloxacin MIC of the *M. tuberculosis* isolate. FRS was rounded to the nearest 0.1 to show the distribution of MIC values at different FRS. Both susceptible and resistant isolates containing alleles encoding the mutation at FRS < 0.9 were included and alleles with FRS >= 0.9 were excluded to avoid overfitting to the majority homogeneous population. R shows the Pearson’s rank correlation coefficient between the unrounded FRS and log_2_ MIC.

## Discussion

We found a near 10% absolute improvement in sensitivity without a significant reduction in specificity when predicting levofloxacin and moxifloxacin resistance using the WHO mutation catalogue by including mixed alleles (Fig. 1c,d). Importantly, this brings the catalogue performance for detecting fluoroquinolone resistance up to the level of isoniazid and rifampicin^4,5^, where the determinants of resistance are also well understood. The magnitude of improvement in sensitivity correlates with the previously estimated 10% frequency of fluoroquinolone resistance conferred by alleles seen in mixed populations^10^. Resistance from minority populations has likewise been implicated for rifampicin, isoniazid and ethambutol^10^, and should be explored further.

Despite the improvement in sensitivity achieved by allowing minor alleles to contribute, 7.9% of levofloxacin resistance and 6.0% of moxifloxacin resistance is not explained by the presence of catalogue mutations. Several scenarios could account for this. Firstly, there are likely additional minor resistance-conferring alleles that could not be detected in this study due to limited sequencing depth (the mean read depth for the CRyPTIC isolates^11^ was 74 ± 44). In addition, our use of two reads in support of an alternative allele to define a minor population is conservative. For example, at a read depth of 30x, a minor allele is only detectable by our approach if at least 6.7% of the population contains it. Secondly, the catalogue mutations are unlikely to be exhaustive as rare resistance conferring mutations would not meet the statistical criteria required^4,5^. Including minor alleles when building catalogues could help improve their comprehensiveness by increasing the number of examples of rarer mutations. Finally, complex resistance mechanisms such as drug efflux may play a role^14^.

The lack of corelation between the magnitude of resistance and the proportion of a population containing a minor resistance conferring allele (Figure 2a-d) is consistent with the suggestion that *gyrA* D94G and A90V do not contribute a significant fitness cost^15^.

Isolates with *gyrA* D94G and A90V seen at very low FRS still had high MICs to levofloxacin and moxifloxacin after two weeks incubation, suggesting that a small resistant population can rapidly outcompete a majority wildtype population under the selective pressure of fluoroquinolone treatment. We infer that it is essential that any tool used for fluoroquinolone DST detects minor resistant populations. This means that for next generation sequencing approaches to be successful, the sequencing depths, variant callers used, and any filters applied need to be carefully optimised, leading to standardised recommendations to prevent the misdiagnosis of fluoroquinolone susceptibility.

Due to the nature of Illumina sequencing performed by the CRyPTIC study, the sequencing depth is limited and highly variable between samples and genetic loci. A deep sequencing approach is necessary to confirm the importance of minority resistant populations in *M. tuberculosis* infection and to better inform the design and parametrisation of bioinformatic pipelines. The Illumina WGS data analysed here provides an estimate of what could be detected by current practice and suggests that high sequencing depths at loci associated with resistance might be required to rule out fluoroquinolone resistance.

To conclude, identifying minority populations containing mutations known to confer resistance improved the sensitivity of fluoroquinolone resistance prediction using the 2021 WHO tuberculosis resistance catalogue. Hence, it is vital that genetics-based tools and pipelines used for fluoroquinolone DST can resolve resistance conferring mutations in minority populations.

## Acknowledgements

We are grateful to the CRyPTIC consortium for helpful discussions and making their data publicly available.

## Funding

A.B. is funded by a NDM Prize Studentship from the Oxford Medical Research Council Doctoral Training Partnership and the Nuffield Department of Clinical Medicine. This work was supported the National Institute for Health Research (NIHR) Oxford Biomedical Research Centre (BRC). The computational aspects of this research were funded from the NIHR Oxford BRC with additional support from a Wellcome Trust Core Award (203141/Z/16/Z). The views expressed are those of the authors and not necessarily those of the NHS, the NIHR or the Department of Health, and the National Institute for Health Research (NIHR) Health Protection Research Unit in Healthcare Associated Infections and Antimicrobial Resistance, a partnership between Public Health England and the University of Oxford, the views expressed are those of the authors and not necessarily those of the NIHR, Public Health England or the Department of Health and Social Care.

## Transparency declarations

The authors have nothing to declare

## Notes

http://ftp.ebi.ac.uk/pub/databases/cryptic/release_june2022/

### Competing Interest Statement

The authors have declared no competing interest.

## References

1 World Health Organization (WHO). Rapid communication: Key changes to the treatment of drug-resistant tuberculosis. (2022).

2 Walker, T. M. et al. Whole-genome sequencing for prediction of Mycobacterium tuberculosis drug susceptibility and resistance: a retrospective cohort study. Lancet Infect Dis 15, 1193–1202, doi:10.1016/S1473-3099(15)00062-6 (2015).

3 Lam, C. et al. Value of routine whole genome sequencing for Mycobacterium tuberculosis drug resistance detection. Int J Infect Dis, doi:10.1016/j.ijid.2021.03.033 (2021).

4 World Health Organization (WHO). Catalogue of mutations in Mycobacterium tuberculosis complex and their association with drug resistance. (2021).

5 Walker, T. M. et al. The 2021 WHO catalogue of Mycobacterium tuberculosis complex mutations associated with drug resistance: a genotypic analysis. The Lancet Microbe 3, e265, doi:10.1016/S2666-5247(21)00301-3 (2022).

6 Nguyen, L. Antibiotic resistance mechanisms in M. tuberculosis: an update. Arch Toxicol 90, 1585–1604, doi:10.1007/s00204-016-1727-6 (2016).

7 Said Mohammed, K. et al. Evaluating the performance of tools used to call minority variants from whole genome short-read data. Wellcome Open Res 3, 21, doi:10.12688/wellcomeopenres.13538.2 (2018).

8 Nimmo, C. et al. Dynamics of within-host Mycobacterium tuberculosis diversity and heteroresistance during treatment. EBioMedicine 55, 102747, doi:10.1016/j.ebiom.2020.102747 (2020).

9 Singhal, R. et al. Sequence Analysis of Fluoroquinolone Resistance-Associated Genes gyrA and gyrB in Clinical Mycobacterium tuberculosis Isolates from Patients Suspected of Having Multidrug-Resistant Tuberculosis in New Delhi, India. J Clin Microbiol 54, 2298–2305, doi:10.1128/JCM.00670-16 (2016).

10 Ye, M. et al. Antibiotic heteroresistance in Mycobacterium tuberculosis isolates: a systematic review and meta-analysis. Ann Clin Microbiol Antimicrob 20, 73, doi:10.1186/s12941-021-00478-z (2021).

11 The CRyPTIC Consortium. A data compendium associating the genomes of 12,289 Mycobacterium tuberculosis isolates with quantitative resistance phenotypes to 13 antibiotics. PLOS Biology 20, e3001721, doi:10.1371/journal.pbio.3001721 (2022).

12 Hunt, M. et al. Minos: variant adjudication and joint genotyping of cohorts of bacterial genomes. bioRxiv, 2021.2009.2015.460475, doi:10.1101/2021.09.15.460475 (2021).

13 The CRyPTIC Consortium. Epidemiological cutoff values for a 96-well broth microdilution plate for high-throughput research antibiotic susceptibility testing of M. tuberculosis. European Respiratory Journal, 2200239, doi:10.1183/13993003.00239-2022 (2022).

14 Remm, S., Earp, J. C., Dick, T., Dartois, V. & Seeger, M. A. Critical discussion on drug efflux in Mycobacterium tuberculosis. FEMS Microbiology Reviews 46, doi:10.1093/femsre/fuab050 (2021).

15 Pi, R., Liu, Q., Takiff, H. E. & Gao, Q. Fitness Cost and Compensatory Evolution in Levofloxacin-Resistant Mycobacterium aurum. Antimicrob Agents Chemother 64, doi:10.1128/aac.00224-20 (2020).

